# Mu and delta opioid receptor antagonists increase the expression of social conditioned place preference in early adolescent mice

**DOI:** 10.1101/2023.07.19.549691

**Authors:** Zofia Harda, Marta Klimczak, Klaudia Misiołek, Magdalena Chrószcz, Aleksandra Rzeszut, Łukasz Szumiec, Maria Kaczmarczyk-Jarosz, Rafał Ryguła, Barbara Ziółkowska, Jan Rodriguez Parkitna

## Abstract

**Rationale:** Social behaviors undergo dramatic changes during adolescence, enabling the development of adult social abilities. These changes are intricately linked to the development of the brain reward system and the activity of endogenous opioid signaling. However, the involvement of the opioid system in the development of social behaviors still raises more questions than answers.

**Objectives:** Here, we investigated the role of the endogenous opioid system in the rewarding effects of social contact in early and late adolescent male mice.

**Methods:** Social reward was assessed using the social conditioned place preference task in early adolescent (~34 days old) and late adolescent (~41 days old) male mice that received a single dose of the selective opioid receptor antagonists cyprodime (1 mg/kg, i.p.), naltrindole (1 mg/kg, i.p.) or norbinaltorphimine (10 mg/kg, i.p.) before the preference posttest.

**Results:** The administration of cyprodime or naltrindole before the posttest significantly increased the preference for the social-conditioned context in early but not late adolescent mice. In contrast, pretreatment with norbinaltorphimine had no effect on context preference.

**Conclusions:** Our findings support a modified version of the state-dependent mu-opioid receptor model of social behavior, where the effects of opioid ligands are not reversed during development but rather weaken or disappear with age. Furthermore, the results indicate that interactions with siblings in early adolescent mice are motivated by negative reinforcement, whereas those in late adolescence are motivated by positive reinforcement.

## Introduction

Affiliative social interactions are usually rewarding. In male and female rodents, social contact with same-sex conspecifics results in a preference for the associated context (Dölen et al., 2013; Nardou et al., 2019; J. B. Panksepp & Lahvis, 2007), and the magnitude of this effect depends on the age and familiarity of the subjects (Harda et al., 2022; Misiołek et al., 2023). The mechanisms underlying the rewarding effects of social interactions have been a focus of intense study, with reported evidence indicating the involvement of the serotonergic, dopaminergic, oxytocin and opioid systems (Borland, 2025; Dölen et al., 2013; Li et al., 2023; Loseth et al., 2014; Meier et al., 2021; Nardou et al., 2019, 2023; Pellissier et al., 2017; Robinson et al., 2002). The role of endogenous opioid signaling is particularly interesting, as it acts as a major modulator of the brain reward system and directly regulates monoaminergic and oxytocin transmission. Notably, drugs that act on opioid receptors have opposite effects on social interactions in rat pups compared with adolescent or adult rodents (Loseth et al., 2014). In particular, opioid agonists decrease maternal contact seeking in isolated rat pups (Gardner, 1985; Kehoe & Blass, 1986), whereas opioid antagonists tend to increase this behavior (Nelson & Panksepp, 1998). In contrast, social play behavior in adolescent rats is facilitated by opioid agonists and attenuated by opioid antagonists (Beatty & Costello, 1982; J. Panksepp et al., 1985; Trezza et al., 2011; L. J. Vanderschuren et al., 1995). Two mechanistic explanations for these contrasting observations have been proposed. The first, proposed by Vanderschuren et al., posits that social play and social contact seeking are qualitatively different behaviors regulated by partially separate neural networks and thus should be considered separately (Guard et al., 2002; L. J. Vanderschuren et al., 1997). Conversely, Loseth and coworkers proposed that the observed age-related differences in the action of opioid receptor ligands stem from developmental changes in response to social isolation, to which animals are usually subjected before social behaviors are tested (Loseth et al., 2014). Infant rodents, which require the presence of caregivers for survival, react to isolation with distress; thus, exogenous mu opioid receptor agonists decrease social contact seeking by reducing negative affect. Older rodents no longer depend on their conspecifics for survival and do not react to isolation with distress. In this state of psychological comfort, social contact is sought for its rewarding effects (Loseth et al., 2014). Since exogenous mu opioid receptor agonists increase the rewarding effects of other types of rewards (MERRER et al., 2009), they also increase the likelihood of seeking social contact in older animals. Notably, the two proposed frameworks are not mutually exclusive and could both be involved in shaping social behaviors. Despite previous attempts to fit the pieces of this puzzle (Herman & Panksepp, 1978; Kennedy et al., 2011), the resulting picture remains ambiguous. Therefore, we employ the social place conditioning paradigm and investigate the effects of selective opioid receptor antagonists to clarify the role of endogenous opioid signaling in social reward among early and late adolescent male mice.

## Methods

### Animals

Experiments were performed on C57BL/6 male mice bred at the Maj Institute of Pharmacology of the Polish Academy of Sciences animal facility. The mice were housed under a 12/12 h light–dark cycle (lights on at 7 AM) under controlled conditions: a temperature of 22 ± 2 °C and a humidity of 40– 60%. After weaning, the mice were housed with littermates of the same sex. Rodent chow and water were available *ad libitum*. Home cages contained aspen or corn bedding. The home and conditioning cages contained aspen nesting material and aspen gnawing blocks. Behavioral tests were conducted during the light phase under dim illumination (5–10 lux). sCPP tests were video recorded with additional infrared LED illumination. All behavioral procedures were approved by the II Local Bioethics Committee in Krakow (permit numbers 35/2019, 265/2019, 305/2020, 32/2021) and performed in accordance with Directive 2010/63/EU of the European Parliament and the Council of 22 September 2010 on the protection of animals used for scientific purposes. The reporting in the manuscript follows the ARRIVE guidelines. Animals were randomly assigned to one of two age groups: early adolescent or middle–late adolescent (referred to as “late adolescent”). The age range was 31–36 (33.8±0.12) days in the posttest for the early adolescent group and 37–47 (41.3±0.16) days for the late adolescent group (a detailed description of the experimental groups is provided in **Table S1**).

### Social conditioned place preference

The test was performed as described previously (Harda et al., 2022) and consisted of three phases: pretest, conditioning, and posttest (**Fig. 1A**). Each cage compartment contained a novel context (context A or context B) defined by the type of bedding and gnawing block size and shape. The bedding types used are listed in **Table S2**, and the materials used are described in **Table S3**. During the pretest, the mice were allowed to explore the test cage freely for 30 minutes, and the time spent in each compartment was recorded. Animals that spent more than 70% of their time in one of the contexts were excluded (**Table S4**). After the pretest, the animals were returned to their home cages for approximately 24 h. Then, the mice were subjected to social conditioning (housing with cage mates) for 24 h in one of the contexts used in the pretest followed by 24 h of isolation conditioning (single housing) in the other context. Conditioning was performed in cages identical to the home cage, with *ad libitum* access to food and water. To prevent bias, the social context was randomly assigned such that approximately half of the animals received social conditioning in context A and half in context B. In cases where the final number of animals conditioned in each context was not equal due to an unequal number of animals in the litter or an unequal number of animals passing the 70% exclusion criterion, randomly selected animals from the larger group were excluded using a Python script (https://zenodo.org/record/8100281). The exclusion was random, but after trimming, the mean 50% context preference during the pretest had to be preserved (for details, see the Supplementary Materials). The conditioning phase lasted 6 days (3 days in each context, alternating every 24 h), and then the posttest was performed in the same way as the pretest. The animals received i.p. one of the following drugs before the posttest: the mu opioid receptor antagonist cyprodime hydrochloride (TOCRIS, cat. no. 2601, 1 mg/kg), the delta opioid receptor antagonist naltrindole hydrochloride (TOCRIS, cat. no. 0740, 1 mg/kg), or the kappa opioid receptor antagonist norbinaltorphimine (TOCRIS, cat. no. 0347 10 mg/kg). Drugs were dissolved in saline and administered intraperitoneally 1 hour (cyprodime and naltrindole) or 4 hours (norbinaltorphimine) before the posttest at a volume of 5 μl/g. The choice of a longer pretreatment time for norbinaltorphimine was dictated by the fact that it is a long-acting antagonist that attains selectivity toward the kappa receptor only by 4 hours after administration, whereas at earlier time points, it also blocks the mu receptor (Endoh et al., 1992). The sCPP results were assessed as the absolute amount of time spent in the social context during the pretest and posttest and as the preference score, which was defined as the difference between the time spent in the social context and the isolation context during the posttest.

**Figure 1.**
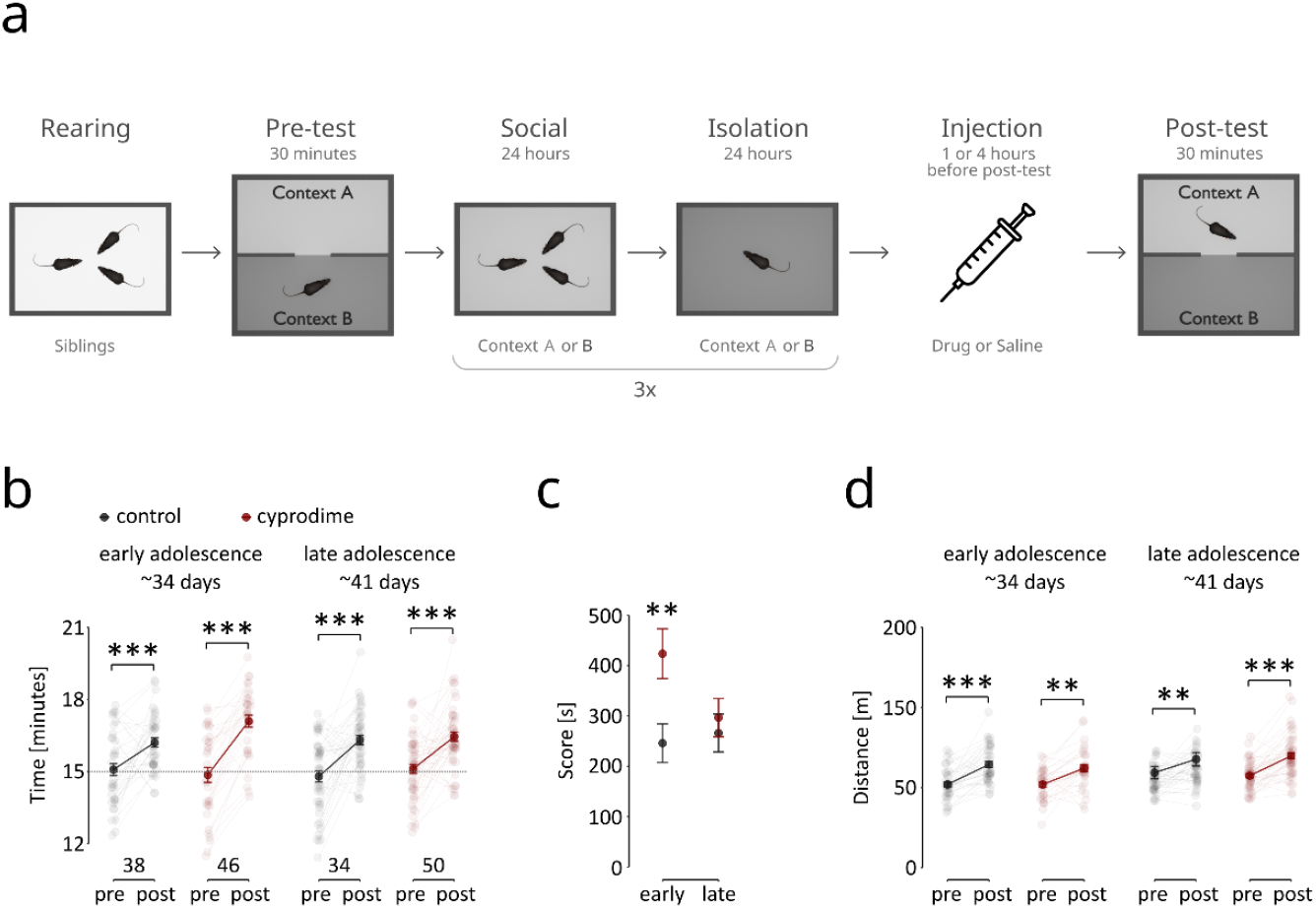
Effect of the selective mu opioid receptor antagonist cyprodime on the expression of sCPP in adolescent male mice. **(a)** A schematic representation of the sCPP procedure. **(b)** The time spent in the social context during the pre- and posttests. The solid circles represent the mean values; black represents the control, saline-treated group, and red represents the cyprodime (1 mg/kg)-treated group. The semitransparent circles represent individual animals, the lines connect the pre- and posttest values for each mouse, and the numbers below indicate group size. The error bars are s.e.m.s. **(c)** The preference score represents the difference in time spent in the social context compared with the isolation context during the posttest. Points represent the mean values ± s.e.m.s, control values are black, and red represents the cyprodime-treated groups. **(d)** Distance moved during the pre- and posttests. The solid points represent the mean values ± s.e.m.s, and the semitransparent points represent individual mice. Two animals had very high locomotor activity (over 200 m traveled) and are not shown on the graph but were included in the analyses. The stars represent significant differences (Tukey HSD) pre vs. post (b, d) or treatment effects (c): ‘^*^’ p <0.05, ‘^**^’ p <0.01, and ‘^***^’ p <0.001.

### Data analysis

The distance traveled and time spent in separate cage compartments were analyzed automatically using EthoVision XT 15 software (Noldus, The Netherlands). Treatment and conditioning effects were assessed using mixed model analysis of variance (type III ANOVA) followed by Tukey’s HSD test. Analyses were performed using R and the implementation of ANOVA in the *car* library with the *afex* addon. Tukey’s HSD was assessed using the *emmeans* library. The statistical significance threshold was set at p < 0.05. The number of mice excluded on the basis of the Grubbs test is listed in **Table S4**. The complete raw behavioral data are deposited in Zenodo: https://doi.org/10.5281/zenodo.14946878.

## Results

The rewarding effects of social interactions were assessed using the social conditioned place preference (sCPP) test in early (~Day 34) and late (~Day 41) adolescent mice (**Fig. 1a**). An age of 36 days was set as the last day of early adolescence based on previous reports on critical adolescence periods for isolation-induced impairments in social and cognitive functioning (Makinodan et al., 2012) and the start of dispersal in *Mus musculus* mice (Groó et al., 2013). The animals received the selective mu opioid receptor antagonist cyprodime (1 mg/kg, i.p.) 1 h before the posttest. Cyprodime significantly increased the time spent in the social context of early adolescent mice but had no effect on late adolescent mice, as confirmed by the significant triple interaction among age, treatment and test phase in ANOVA (**Fig. 1b**, three-way ANOVA: Pre-Post F_1,164_ = 107.64, p < 0.0001; Treatment F_1,164_ = 2.85, p = 0.093; Treatment × Pre-Post F_1,164_ = 2.43, p = 0.121; Treatment × Pre-Post × Age F_1,164_ = 4.66, p = 0.032; all other contrasts F < 1). Among the early adolescent mice (P34), 68% in the control group and 94% in the cyprodime group spent more time in the social context from the pretest to the posttest. These numbers were similar in the control and cyprodime-treated groups (74% and 78%, respectively). Cyprodime administration also significantly increased the preference score in early (but not late) adolescent mice. (**Fig. 1c**, two-way ANOVA: Treatment F_1,164_ = 6.47, p = 0.012; Age F_1,164_ = 1.7028, p = 0.194; Treatment × Age F_1,164_ = 3.2280, p = 0.074). There was no effect of treatment on locomotor activity, although there was a significant increase in activity with age and between the pre- and posttests (**Fig. 1d;** three-way ANOVA: Pre-Post F_1,164_ = 63.44, p < 0.0001; Age F_1,164_ = 6.86, p = 0.010; Treatment × Pre-Post × Age F_1,164_=1.34, p = 0.249; all other contrasts F < 1). These data show that mu opioid receptors play an important role in the regulation of social reward in early adolescence but not in late adolescence.

A similar effect was observed after treatment with naltrindole (1 mg/kg, i.p., 1 h prior to the posttest). There was a significant triple interaction between treatment, age, and test phase (**Fig. 2a**, three-way ANOVA: Pre-Post F_1,108_ = 34.10, p < 0.0001; Treatment × Age F_1,108_ = 1.15, p = 0.286; Treatment × Pre-Post × Age F_1,108_ = 4.26, p = 0.041; all other contrasts F < 1). The fraction of early adolescent animals that exhibited an increase in the time spent in the social context after conditioning was slightly greater in the drug-treated group than in the saline group (saline: 68%, naltrindole: 73%), whereas for late adolescent animals, this pattern was reversed (saline: 76%, naltrindole: 63%). There was a significant interaction effect between naltrindole treatment and age in the preference score, with drug-treated early adolescent animals showing a greater preference for the conditioned context (**Fig. 2b**, two-way ANOVA: Treatment F_1,108_ = 1.21, p = 0.274; Age F_1,108_ = 1.03, p = 0.313; Treatment × Age F_1,108_ = 5.77, p = 0.018). Locomotor activity was not appreciably affected by treatment, although it clearly increased during the posttest compared with the pretest (**Fig. 2c**, three-way ANOVA: Pre-Post F_1,108_ = 92.15, p < 0.0001; Age F_1,108_ = 2.29, p = 0.133; Treatment × Pre-Post F_1,108_ = 1.91, p = 0.170; Treatment × Pre-Post × Age F_1,108_ = 1.21, p = 0.274; all other contrasts F < 1). These data show that delta opioid receptors differentially regulate social reward in early and late adolescence.

**Figure 2.**
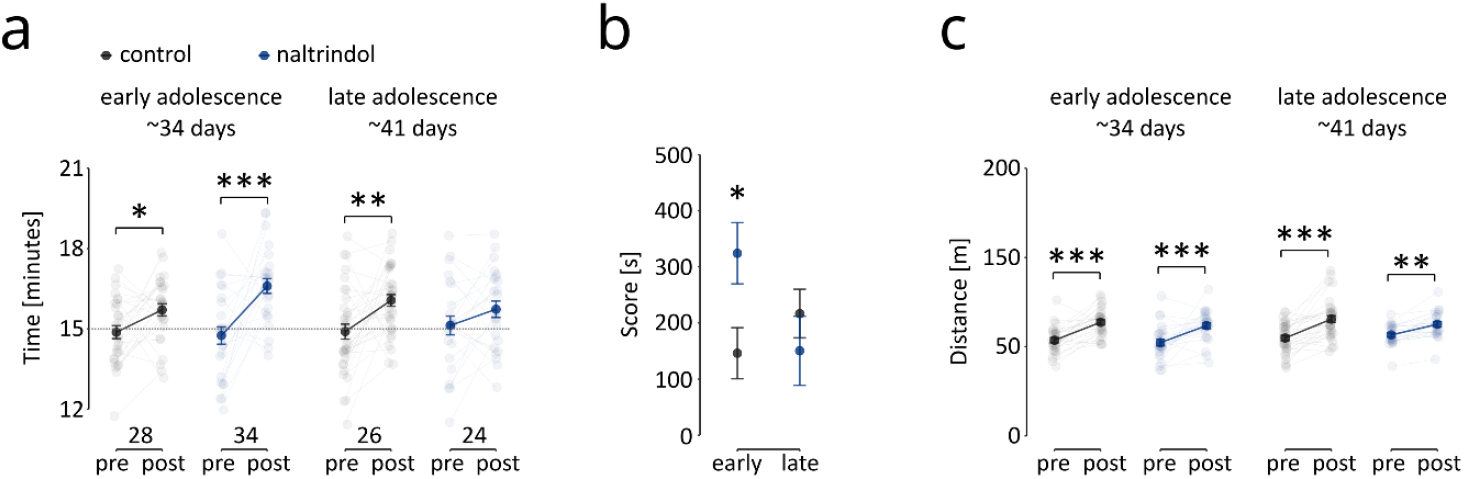
Effect of the selective delta opioid receptor antagonist naltrindole on sCPP in adolescent male mice. **(a)** Time spent in the social context during the pre- and posttests. The solid circles represent the mean values, the black circles represent the control, and the blue circles represent the naltrindole (1 mg/kg)-treated group. Semitransparent circles represent individual animals, lines connect pre- and posttest values for individual mice, and the numbers below indicate group size. The error bars are s.e.m.s. **(b)** sCPP preference score. Points represent the mean values ± s.e.m.s, control values are black, and blue represents the naltrindole-treated groups. **(c)** Distance moved during the pre- and posttests. The solid points represent the mean values ± s.e.m.s, and the semitransparent points represent individual mice. The stars represent significant differences (Tukey HSD) pre vs. post-test (a, c) or treatment effects (b): ‘^*^’ p <0.05, and ‘^***^’ p <0.001.

Finally, the kappa opioid receptor antagonist norbinaltorphimine (10 mg/kg, i.p., 4 h prior to posttest) had no significant effect on the time spent in the social context (**Fig. 3a**, three-way ANOVA: Pre-Post F_1,76_ = 36.78, p < 0.0001; Age F_1,76_ = 2.98, p < 0.088; Pre-Post × Age F_1,76_ = 3.68, p = 0.059; Treatment × Pre-Post × Age F_1,76_ = 0.96, p = 0.330; all other contrasts F < 1). The percentages of early adolescent animals that spent more time in the social context after conditioning were 63% and 44% in the saline and norbinaltorphimine groups, respectively. In the case of late adolescent animals, the percentages of animals that exhibited an increase in the time spent in the social context after conditioning were 78% and 75% for the saline and norbinaltorphimine treatment groups, respectively. Accordingly, norbinaltorphimine had no significant effect on context preference on the basis of score values, although there was a significant, treatment-independent effect of age, with higher scores observed in late adolescent animals (**Fig. 3b**, two-way ANOVA: Treatment F_1,76_ = 0.903, p = 0.346; Age F_1,76_ = 6.302, p = 0.014; Treatment × Age F_1,76_ = 0.070, p = 0.792). Treatment had no effect on locomotor activity, although again, there was significantly greater activity in the posttest, higher activity in late adolescent mice, and a significant interaction between these two factors (**Fig. 3c**, three-way ANOVA: Pre-Post F_1,76_= 36.78, p < 0.0001; Age F_1,76_ = 11.12, p = 0.001; Pre-Post × Age F_1,76_ = 6.28, p = 0.014; Treatment × Pre-Post F_1,76_ = 1.24, p = 0.268; all other contrasts F < 1). These data show that kappa opioid receptors do not play a crucial role in the regulation of social reward in adolescent mice.

**Figure 3.**
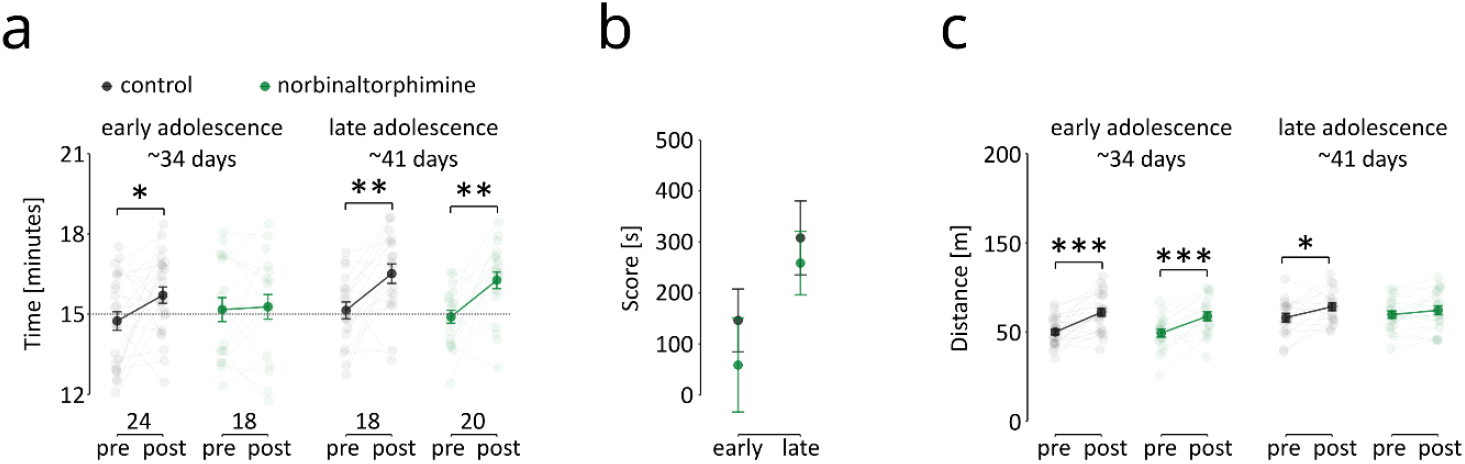
Effect of the kappa opioid receptor antagonist norbinaltorphimine on sCPP in adolescent male mice. **(a)** Time spent in the social context during the pre- and posttests. The solid circles represent the mean values, black represents the control, and green represents the norbinaltorphimine (10 mg/kg)-treated group. Semitransparent circles represent individual animals, lines connect pre- and posttest values for individual mice, and the numbers below indicate group size.. The error bars are s.e.m.s. **(b)** sCPP preference score. Points represent the mean values ± s.e.m.s, control values are black, and green represents the norbinaltorphimine-treated groups. **(c)** Distance moved during the pre- and posttests. The solid points represent the mean values ± s.e.m.s, and the semitransparent points represent individual mice. The stars represent significant differences (Tukey HSD) pre vs. post-test (a, c): ‘^*^’ p <0.05, ‘^**^’ p <0.01, and ‘^***^’ p <0.001.

## Discussion

Here, we show that the selective mu opioid receptor antagonist cyprodime and the delta opioid receptor antagonist naltrindole increased the expression of sCPP in early, but not late, adolescent mice, whereas a kappa opioid receptor antagonist (norbinaltorphimine) had no significant effect. Notably, the effects of antagonists probably result from blocking endogenous opioid signaling, which strongly implicates enkephalins, the primary mu and delta receptor agonists, as regulators of the expression of sCPPs. Moreover, we note that none of the antagonists used here were previously observed to have rewarding or aversive properties in the conditioned place preference task (Sikora et al., 2019); thus, social preference is not confounded by a change in the affective state of the animals. Finally, in some of the experiments, there was an increase in preference for the social context in late adolescent vs. early adolescent mice; however, this effect was not consistent and requires further investigation.

Two studies to date have investigated the effects of opioid ligands on social behavior after a period of social isolation in prepubertal vs. postpubertal or adult rodents (Herman & Panksepp, 1978; Kennedy et al., 2011). Kennedy and collaborators studied the effect of morphine on social interaction with a cagemate in mice. Morphine decreased the time spent investigating conspecifics in late juvenile, late adolescent and adult mice, but sensitivity to the drug decreased with age. In the second study, Herman and Panksepp investigated the effects of naloxone on separation-induced distress vocalizations in juvenile and adult guinea pigs and reported an increase in this measure of behavior in both age groups. However, as noted by the authors, the interpretation of these results is most likely confounded by the ceiling effect because the baseline level of distress vocalizations is very high in pups. Our findings, together with those of Kennedy and colleagues (Kennedy et al., 2011), suggest that a modified version of the state-dependent mu opioid receptor model of social behavior (Loseth et al., 2014) could explain the developmental differences in the effects of mu opioid receptor ligands on social interactions. The model presented by Loseth and coworkers is based on observations that opioid ligands have opposite effects on social contact seeking in neonates and social play behavior in adolescent rodents. Conversely, the data presented here suggest that the effects of opioid ligands do not switch direction at some point in development. Instead, their effects probably weaken or disappear with age.

An alternative explanation for the paradoxical actions of mu opioid receptor ligands, proposed by Vanderschuren and colleagues, is that social contact seeking and social play differ qualitatively (L. J. Vanderschuren et al., 1997). This hypothesis was based on observations that mu opioid agonists increase social play behavior but decrease social contact time in the same individuals (L. J. M. J. Vanderschuren et al., 1995). Several other observations may support this hypothesis. It has been reported that mu opioid receptor agonists, including morphine, beta-endorphin, methadone, and DAMGO, decrease social contact time in adolescent (Hol et al., 1996; Hughes et al., 2021; Niesink & Van Ree, 1982; J. Panksepp et al., 1979) and adult rodents (Deak et al., 2009; Kudryavtseva et al., 2004; Meyerson, 1981; J. Panksepp et al., 1979; Piccin et al., 2022; Plonsky & Freeman, 1982; Šlamberová et al., 2016), either housed in groups (Meyerson, 1981; J. Panksepp et al., 1979; Piccin et al., 2022; Šlamberová et al., 2016) or isolated (Deak et al., 2009; Hol et al., 1996; Hughes et al., 2021; Kudryavtseva et al., 2004; Niesink & Van Ree, 1982; Plonsky & Freeman, 1982). Together, these reports indicate that the activation of opioid receptors decreases social contact seeking irrespective of age or potential isolation effects. On the other hand, treatment with morphine increases social play behavior in juvenile rats, and mu opioid receptor antagonists decrease this behavior, as does place preference conditioned by exposure to a play partner (Beatty & Costello, 1982; J. Panksepp et al., 1985; Trezza et al., 2011; L. J. Vanderschuren et al., 1995). These results indicate that social interaction and social play indeed represent distinct aspects of behavior, regulated by the mu opioid system in the opposite way (Guard et al., 2002; L. J. Vanderschuren et al., 1997). Together, our data and earlier studies suggest that both hypotheses presented in the introduction are valid. The model proposed by Loseth and colleagues and extended here could explain developmental changes in opioid receptor function, whereas the hypothesis presented by Vanderschuren and colleagues offers a plausible explanation for the differences in the influence of opioid ligands on social play versus social contact seeking.

Finally, it is important to consider the results presented here in the context of the framework proposed by Riters and collaborators concerning the involvement of opioid receptors in gregarious social interactions (Riters et al., 2019). This theory is based on the notion that joining a social group might be motivated by both negative and positive reinforcement. Specifically, grouping might be strengthened because it reduces the negative affect caused by social isolation or because interactions with conspecifics are rewarding. Since mu opioid agonists decrease, while antagonists increase, the effects of aversive conditioning (Meier et al., 2021), our findings suggest that interactions with siblings in early adolescent mice are motivated by negative reinforcement, whereas those in late adolescent mice are motivated by positive reinforcement.

Taken together, our results confirm the presence of distinct mechanisms controlling social behaviors during development and show that mu- and delta-opioid receptors play major roles in controlling social contact-conditioned responses in early adolescent but not late adolescent male mice.

## Abbreviations

sCPP: social conditioned place preference

## Acknowledgements

We would like to thank Dr. Jakub Dzik from the Nencki Institute for Experimental Biology, Polish Academy of Sciences, for the Python script used for social conditioned place preference data trimming (https://zenodo.org/record/8100281).

## Ethical Approval

All behavioral procedures were approved by the II Local Bioethics Committee in Krakow (permit numbers 35/2019, 265/2019, 305/2020, 32/2021) and performed in accordance with Directive 2010/63/EU of the European Parliament and the Council of 22 September 2010 on the protection of animals used for scientific purposes.

## Consent to participate

Not applicable.

## Consent to publish

All Authors consented to publication.

## Data Availability Statement

The complete raw behavioral data are deposited in Zenodo: https://doi.org/10.5281/zenodo.14946878.

## Authors Contributions

**ZH, RR, BZ** and **JRP** designed the study with input from all the Authors; **ZH, MK, KM, MC, AR, ŁS** and **MK-J** performed the experiments; **ZH** and **JRP** analyzed the data; **ZH, RR, BZ** and **JRP** wrote the manuscript with help from all the Authors.

## References

Beatty, W. W., & Costello, K. B. (1982). Naloxone and play fighting in juvenile rats. Pharmacology Biochemistry and Behavior, 17(5), 905–907. 10.1016/0091-3057(82)90470-1

Borland, J. M. (2025). A review of the effects of different types of social behaviors on the recruitment of neuropeptides and neurotransmitters in the nucleus accumbens. Frontiers in Neuroendocrinology, 77, 101175. 10.1016/j.yfrne.2025.101175

Deak, T., Arakawa, H., Bekkedal, M., & Panksepp, J. (2009). Validation of a novel social investigation task that may dissociate social motivation from exploratory activity— ScienceDirect. Behavioural Brain Research, 16;199(2), 326–333.

Dölen, G., Darvishzadeh, A., Huang, K. W., & Malenka, R. C. (2013). Social reward requires coordinated activity of accumbens oxytocin and 5HT. Nature, 501(7466), 179–184. 10.1038/nature12518

Endoh, T., Matsuura, H., Tanaka, C., & Nagase, H. (1992). Nor-binaltorphimine: A potent and selective kappa-opioid receptor antagonist with long-lasting activity in vivo. Archives Internationales De Pharmacodynamie Et De Therapie, 316, 30–42.

Gardner, C. R. (1985). Distress vocalization in rat pups a simple screening method for anxiolytic drugs. Journal of Pharmacological Methods, 14(3), 181–187. 10.1016/0160-5402(85)90031-2

Guard, H. J., Newman, J. D., & Roberts, R. L. (2002). Morphine administration selectively facilitates social play in common marmosets. Developmental Psychobiology, 41(1), 37–49. 10.1002/dev.10043

Harda, Z., Chrószcz, M., Misiołek, K., Klimczak, M., Szumiec, Ł., Kaczmarczyk-Jarosz, M., & Rodriguez Parkitna, J. (2022). Establishment of a social conditioned place preference paradigm for the study of social reward in female mice. Scientific Reports, 12(1), 11271. 10.1038/s41598-022-15427-9

Herman, B. H., & Panksepp, J. (1978). Effects of morphine and naloxone on separation distress and approach attachment: Evidence for opiate mediation of social affect. Pharmacology Biochemistry and Behavior, 9(2), 213–220. 10.1016/0091-3057(78)90167-3

Hol, T., Ruven, S., Van Ree, J. M., & Spruijt, B. M. (1996). Chronic administration of Org2766 and morphine counteracts isolation-induced increase in social interest: Implication of endogenous opioid systems. Neuropeptides, 30(3), 283–291. 10.1016/S0143-4179(96)90074-8

Hughes, E. M., Calcagno, P., Sanchez, C., Smith, K., Kelly, J. P., Finn, D. P., & Roche, M. (2021). Mu-opioid receptor agonism differentially alters social behaviour and immediate early gene expression in male adolescent rats prenatally exposed to valproic acid versus controls. Brain Research Bulletin, 174, 260–267. 10.1016/j.brainresbull.2021.06.018

Kehoe, P., & Blass, E. M. (1986). Opioid-mediation of separation distress in 10-day-old rats: Reversal of stress with maternal stimuli. Developmental Psychobiology, 19(4), 385–398. 10.1002/dev.420190410

Kennedy, B. C., Panksepp, J. B., Wong, J. C., Krause, E. J., & Lahvis, G. P. (2011). Age-dependent and strain-dependent influences of morphine on mouse social investigation behavior. Behavioural Pharmacology, 22(2), 147. 10.1097/FBP.0b013e328343d7dd

Kudryavtseva, N. N., Gerrits, M. A. F. M., Avgustinovich, D. F., Tenditnik, M. V., & Van Ree, J. M. (2004). Modulation of anxiety-related behaviors by mu- and kappa-opioid receptor agonists depends on the social status of mice. Peptides, 25(8), 1355–1363. 10.1016/j.peptides.2004.05.005

Li, X., Wu, J., Li, X., & Zhang, J. (2023). The effect of intraperitoneal and intra-RMTg infusions of CTAP on rats’ social interaction. Behavioural Brain Research, 446, 114333. 10.1016/j.bbr.2023.114333

Loseth, G. E., Ellingsen, D.-M., & Leknes, S. (2014). State-dependent μ-opioid modulation of social motivation. Frontiers in Behavioral Neuroscience, 8, 430. 10.3389/fnbeh.2014.00430

Meier, I. M., van Honk, J., Bos, P. A., & Terburg, D. (2021). A mu-opioid feedback model of human social behavior. Neuroscience & Biobehavioral Reviews, 121, 250–258. 10.1016/j.neubiorev.2020.12.013

Merrer, J. L., Becker, J. A. J., Befort, K., & Kieffer, B. L. (2009). Reward Processing by the Opioid System in the Brain. Physiological Reviews, 89(4), 1379–1412. 10.1152/physrev.00005.2009

Meyerson, B. J. (1981). Comparison of the effects of β-endorphin and morphine on exploratory and socio-sexual behaviour in the male rat. European Journal of Pharmacology, 69(4), 453–463. 10.1016/0014-2999(81)90449-0

Misiołek, K., Klimczak, M., Chrószcz, M., Szumiec, Ł., Bryksa, A., Przyborowicz, K., Rodriguez Parkitna, J., & Harda, Z. (2023). Prosocial behavior, social reward and affective state discrimination in adult male and female mice. Scientific Reports, 13(1), Article 1. 10.1038/s41598-023-32682-6

Nardou, R., Lewis, E. M., Rothhaas, R., Xu, R., Yang, A., Boyden, E., & Dölen, G. (2019). Oxytocin-dependent reopening of a social reward learning critical period with MDMA. Nature, 569(7754), 116–120. 10.1038/s41586-019-1075-9

Nardou, R., Sawyer, E., Song, Y. J., Wilkinson, M., Padovan-Hernandez, Y., de Deus, J. L., Wright, N., Lama, C., Faltin, S., Goff, L. A., Stein-O’Brien, G. L., & Dölen, G. (2023). Psychedelics reopen the social reward learning critical period. Nature, 618(7966), 790–798. 10.1038/s41586-023-06204-3

Nelson, E. E., & Panksepp, J. (1998). Brain substrates of infant-mother attachment: Contributions of opioids, oxytocin, and norepinephrine. Neuroscience and Biobehavioral Reviews, 22(3), 437–452. 10.1016/s0149-7634(97)00052-3

Niesink, R. J. M., & Van Ree, J. M. (1982). Antidepressant drugs normalize the increased social behaviour of pairs of male rats induced by short term isolation. Neuropharmacology, 21(12), 1343–1348. 10.1016/0028-3908(82)90144-7

Panksepp, J. B., & Lahvis, G. P. (2007). Social reward among juvenile mice. Genes, Brain, and Behavior, 6(7), 661–671. 10.1111/j.1601-183X.2006.00295.x

Panksepp, J., Jalowiec, J., DeEskinazi, F. G., & Bishop, P. (1985). Opiates and play dominance in juvenile rats. Behavioral Neuroscience, 99(3), 441–453. 10.1037//0735-7044.99.3.441

Panksepp, J., Najam, N., & Soares, F. (1979). Morphine reduces social cohesion in rats. Pharmacology, Biochemistry, and Behavior, 11(2), 131–134. 10.1016/0091-3057(79)90002-9

Pellissier, L. P., Gandía, J., Laboute, T., Becker, J. A. J., & Le Merrer, J. (2017). μ opioid receptor, social behaviour and autism spectrum disorder: Reward matters. British Journal of Pharmacology. 10.1111/bph.13808

Piccin, A., Courtand, G., & Contarino, A. (2022). Morphine reduces the interest for natural rewards. Psychopharmacology, 239(8), 2407–2419. 10.1007/s00213-022-06131-7

Plonsky, M., & Freeman, P. R. (1982). The effects of methadone on the social behavior and activity of the rat. Pharmacology Biochemistry and Behavior, 16(4), 569–571. 10.1016/0091-3057(82)90417-8

Riters, L. V., Kelm-Nelson, C. A., & Spool, J. A. (2019). Why Do Birds Flock? A Role for Opioids in the Reinforcement of Gregarious Social Interactions. Frontiers in Physiology, 10, 421. 10.3389/fphys.2019.00421

Robinson, D. L., Heien, M. L. A. V., & Wightman, R. M. (2002). Frequency of dopamine concentration transients increases in dorsal and ventral striatum of male rats during introduction of conspecifics. The Journal of Neuroscience: The Official Journal of the Society for Neuroscience, 22(23), 10477–10486. 10.1523/JNEUROSCI.22-23-10477.2002

Sikora, M., Skupio, U., Jastrzebska, K., Rodriguez Parkitna, J., & Przewlocki, R. (2019). Antagonism of μ-opioid receptors reduces sensation seeking-like behavior in mice. Behavioural Brain Research, 359, 498–501. 10.1016/j.bbr.2018.11.039

Šlamberová, R., Mikulecká, A., Macúchová, E., Hrebíčková, I., Ševčíková, M., Nohejlová, K., & Pometlová, M. (2016). Morphine Decreases Social Interaction of Adult Male Rats, While THC Does Not Affect It. Physiological Research, S547–S555. 10.33549/physiolres.933527

Trezza, V., Damsteegt, R., Achterberg, E. J. M., & Vanderschuren, L. J. M. J. (2011). Nucleus Accumbens μ-Opioid Receptors Mediate Social Reward. Journal of Neuroscience, 31(17), 6362–6370. 10.1523/JNEUROSCI.5492-10.2011

Vanderschuren, L. J. M. J., Spruijt, B. M., Van Ree, J. M., & Niesink, R. J. M. (1995). Effects of morphine on different aspects of social play in juvenile rats. Psychopharmacology, 117(2), 225–231. 10.1007/BF02245191

Vanderschuren, L. J., Niesink, R. J., Spruijt, B. M., & Van Ree, J. M. (1995). Mu- and kappa-opioid receptor-mediated opioid effects on social play in juvenile rats. European Journal of Pharmacology, 276(3), 257–266. 10.1016/0014-2999(95)00040-r

Vanderschuren, L. J., Niesink, R. J., & Van Ree, J. M. (1997). The neurobiology of social play behavior in rats. Neuroscience and Biobehavioral Reviews, 21(3), 309–326. 10.1016/s0149-7634(96)00020-6

